# 5-MeO-DMT modifies innate behaviors and promotes structural neural plasticity in mice

**DOI:** 10.1101/2022.11.03.515044

**Authors:** Sarah J. Jefferson, Ian Gregg, Mark Dibbs, Clara Liao, Hao Wu, Pasha A. Davoudian, Jeffrey S. Sprouse, Alexander M. Sherwood, Alfred P. Kaye, Christopher Pittenger, Alex C. Kwan

## Abstract

Serotonergic psychedelics are gaining increasing interest as potential therapeutics for a range of mental illnesses. Compounds with short-lived subjective effects may be clinically useful because dosing time would be reduced, which may improve patient access. One short-acting psychedelic is 5-MeO-DMT, which has been associated with improvement in depression and anxiety symptoms in early clinical studies. However relatively little is known about the behavioral effects and neural mechanisms of 5-MeO-DMT in animal models. Here we characterized the effects of 5-MeO-DMT on innate behaviors and dendritic architecture in mice. We showed that 5-MeO-DMT induces a dose-dependent increase in head-twitch response that is shorter in duration than that induced by psilocybin at all doses tested. 5-MeO-DMT also substantially suppresses social ultrasonic vocalizations produced during mating behavior. 5-MeO-DMT produces long-lasting increases in dendritic spine density in the mouse medial frontal cortex that are driven by an elevated rate of spine formation. However, unlike psilocybin, 5-MeO-DMT did not affect the size of dendritic spines. These data provide insights into the behavioral and neural consequences underlying the action of 5-MeO-DMT and highlight similarities and differences with those of psilocybin.

## INTRODUCTION

Psychedelics are emerging as a promising potential treatment for mental illnesses [1-5]. Specifically, psilocybin-assisted psychotherapy has been shown to be efficacious for major depression and treatment-resistant depression in Phase 2 clinical trials [6,7]. The most notable outcome from these trials is the enduring nature of the beneficial effect: after one or two dosing sessions, reduction of depressive symptoms could still be observed after 4 weeks [7] or, in another case, after 3 months [8]. However, psilocybin acutely alters the states of perception and cognition for 2 – 3.5 hr in humans [9], and therefore dosing requires close supervision of medical professionals, which may be expensive and can reduce patient access. There is growing interest in other psychedelics and routes of administration that produce a shorter duration of action, such as 5-methoxy-*N,N*-dimethyltryptamine (5-MeO-DMT), which has a rapid onset of action – within 1 minute – and is psychoactive for only <20 min after inhalation [10-12]. Despite the more transient action, intensities of the peak drug-evoked psychological experience are comparable for 5-MeO-DMT and psilocybin [13]. 5-MeO-DMT has been found to be generally safe with few adverse effects [10,12]. Moreover, open-label studies suggest that the use of 5-MeO-DMT is associated with improvements in depression and anxiety [14,15]. Therefore, there are preliminary but encouraging clues to suggest that 5-MeO-DMT may be useful for treating psychiatric disorders, with an advantage of brief acute action that is more compatible with use in the clinic.

5-MeO-DMT, a compound found naturally in the Sonoran Desert toad, has been noted for its unusual pharmacological properties and clinical significance (reviewed in [16,17]). As a tryptamine psychedelic like psilocybin, 5-MeO-DMT targets a similar set of serotonin receptors including the 5-HT_1A_ and 5-HT_2A_ subtypes [18,19], although 5-MeO-DMT has a higher affinity preference for 5-HT_1A_ receptors, relative to 5-HT_2A_, receptors than psilocybin [20]. It has been argued that this balance of 5-HT_1A_ versus 5-HT_2A_ receptor agonism may be important for the effects of psychedelics [21,22]. Researchers have characterized the dose-response curve for 5-MeO-DMT using head-twitch response [23,24] and drug discrimination assays [18]. Early studies indicate that, like other serotonergic psychedelics, 5-MeO-DMT reliably suppress firing activity in dorsal raphe [25,26]. There are reports that 5-MeO-DMT may decrease anxiety-like behavior in mice [27], enhance cell proliferation in dentate gyrus in mice [28], alter expression of inflammatory and LTP-associated proteins in human brain organoids [29], and perturb prefrontal cortical activity in rats [30]. In cultured cortical neurons, incubation with 5-MeO-DMT increases dendritic arbor complexity, suggesting a capacity to enhance neural plasticity [24].

Whether 5-MeO-DMT can evoke neural plasticity *in vivo* is not known. For psilocybin, in a prior study [31], we showed that a single dose increases the density of dendritic spines in the mouse medial frontal cortex. When tracking the same spines with chronic two-photon microscopy over time, we found that the elevated spine density persists for 1 month, paralleling the long-lasting behavioral effects reported in humans [31]. These plasticity effects are consistent with studies *in vitro* [32,33], as well in mouse hippocampus and in pigs [34,35] and similar studies of related compounds [36,37].

It is unknown whether 5-MeO-DMT has similar effects on plasticity; given that its acute actions are shorter, its neural effects may differ from those of psilocybin. Are the durations of acute and long-lasting effects related, which would predict that a shorter-acting psychedelic like 5-MeO-DMT would yield shorter-lasting effects on neural plasticity? Alternatively, does 5-MeO-DMT evoke comparable long-lasting effects on neuronal architecture?

To address this question, we studied the effects of 5-MeO-DMT in mice, pairing our study on longitudinal neural plasticity with relevant assays of innate behavior. We tested the behavioral consequences using two assays of innate behavior: head-twitch response and social ultrasonic vocalization (USV). The results show that, consistent with human data, 5-MeO-DMT elicits acute behavioral effects that are more transient than those of psilocybin. We then measured the impact of 5-MeO-DMT on dendritic architecture, using longitudinal *in vivo* two-photon microscopy. We found that 5-MeO-DMT, like psilocybin, produces a prolonged increase in dendritic spine density. However, unlike psilocybin, 5-MeO-DMT did not increase the dendritic spine size. Together, these results provide insights into the behavioral effects and neural mechanisms underlying the action of 5-MeO-DMT, highlighting both similarities to and differences from psilocybin.

## MATERIALS AND METHODS

### Animals

Experiments were performed on males and females, except for the USV measurements, because males vocalize substantially more than females. Animals were randomly assigned to treatment groups. For behavioral experiments, C57BL/6J mice were used (Stock #000664, Jackson Laboratory). For imaging experiments, *Thy1*^*GFP*^ line M mice were used (Tg(Thy1-EGFP)MJrs/J, Stock #007788, Jackson Laboratory). Animals were group-housed (2 – 5 mice per cage) in a facility with 12 hr light-dark cycle (7:00 AM – 7:00 PM for light) with ad libitum access to food and water. Experimental procedures were approved by the Institutional Animal Care and Use Committee at Yale University.

### Drugs

Ketamine working solution (1 mg/mL) was prepared by diluting a stock solution (Ketathesia 100 mg/mL, 10mL; #55853, Henry Schein) with sterile saline (0.9% sodium chloride, Hospira). Psilocybin working solution (0.025 – 0.8 mg/mL) was prepared by dissolving powder obtained from Usona Institute’s Investigational Drug & Material Supply Program with saline. 5-MeO-DMT was obtained from two sources. Initially, we obtained 5-MeO-DMT in freebase form (#11480, Cayman Chemical). A 7 mg/mL stock solution was prepared by dissolving 5-MeO-DMT powder in 50 μL ethanol followed by 950 μL saline. 5-MeO-DMT working solution (2 mg/mL) was prepared by further diluting the 7mg /mL stock solution with saline. Freebase 5-MeO-DMT was used for the first set of longitudinal two-photon imaging (n = 2 mice) with the appropriate vehicle control (1.4% ethanol in saline, n = 2 mice), and for ultrasonic vocalization measurements with psilocybin, ketamine, and saline for comparison. Later, we obtained 5-MeO-DMT in salt form, i.e., 5-MeO-DMT succinate [38], from Usona Institute’s Investigational Drug & Material Supply Program. In this case, a working solution was prepared by diluting 3.08 mg (equivalent to 2 mg freebase) of the powder into 1 mL saline. 5-MeO-DMT succinate was used for the second set of longitudinal two-photon imaging (n = 4 mice) with saline as vehicle control (n = 2 mice), and for head-twitch response measurements. We note that 5-MeO-DMT in solution, regardless of whether made from freebase or salt, rapidly oxidizes and loses potency. Therefore we prepared fresh solution every day for the experiments. Ketamine working solution and psilocybin stock and working solutions were replaced approximately every 30 days.

### Head-twitch response

To detect head movement, we used magnetic ear tags. Our scheme followed earlier work that have recorded head-twitch responses using magnets based on the same principle of electromagnetic induction [39,40]. An ear tag was made by affixing a neodymium magnet (N45, diameter: 3 mm diameter, 0.5 mm thick, #D1005-10, SuperMagnetMan) to an aluminum ear tag (La Pias #56780, Stoelting) using cyanoacrylate (Super Glue Ultra Gel Control, #1739050, Loctite). The magnet was coated with a nitrocellulose marker (#7056, ColorTone) and dried for 2-4 hours to dry to prevent the magnet from irritating the mouse’s ear. The mouse was briefly anesthetized with isofluorane and the magnetic ear tag was placed on a mouse’s ear using the ear tag applicator (#56791 Stoelting). Each mouse received one ear tag because signal was sufficiently strong with one tag and ears tended to get stuck together when both received ear tags. After attaching the ear tags, mice recovered for at least 2 days before testing. To record head-twitch responses, the mouse was placed inside a clear plastic, open-top container (4” x 4” x 4” height). An 820-ft-long, 30 AWG enameled copper wire was wound around the container, with its two ends connected to a preamplifier (PP444, Pyle), which was recorded via a data acquisition device (USB-6001, National Instruments) using MATLAB in a desktop computer. Head movements would induce a changing magnetic flux within the coil loop, resulting in a voltage change that can be measured. For each mouse, we recorded for 120 min immediately after intraperitoneal injection of drug or vehicle. We applied a 70-110 Hz bandpass filter to the recorded signal to isolate the ∼90 Hz head-twitch from other head movements, and then using a peak detection algorithm to detect individual head-twitch events. For analysis, we calculated total count, which is the number of head-twitch responses detected over the entire recording, and duration, which is the full width at half maximum with maximum defined as the peak of the head-twitch response rate. In a subset of experiments, we simultaneously recorded the animal using a high-speed camera setup described previously [31]. This allowed us to compare head-twitch responses detected automatically via magnetic ear tags versus manually via visual inspection of videos to determine the performance of the magnetic ear tag system. The parts list, step-by-step instructions, and software for building a magnetic ear tag reporter setup for measuring head-twitch response are available at https://github.com/Kwan-Lab/HTR.

### Social ultrasonic vocalization (USV)

To obtain consistent social USVs in male mice, we followed a habituation protocol described previously [41]. C57BL/6J (Stock No. 000664) mice were bred in-house with the resulting pups being used for experiments after weaning and socialization. Socialization occurred from the time of weaning until 50 days after birth and involved putting a male-female pair of juvenile mice into a cylindrical recording chamber (6” diameter x 7” height, acrylic exterior with rubber dampening mat) within a dark, soundproof outer box and allowing them to interact for 2 to 3 minutes. Vocalizations were recorded using a condenser ultrasound microphone (CM16/CMPA, Avisoft Bioacoustics) connected to a recording interface unit (UltraSoundGate 116Hb, Avisoft Bioacoustics). The Avisoft RECORDER software was used to generate audio files in .wav format. Socialization took place 3 times a week, for 3 consecutive weeks, until the male mice were vocalizing robustly. Male mice that failed to produce loud, consistent vocalizations over multiple consecutive sessions were removed from the study. The remaining male mice were put through three iterations of a 4-day vocalization protocol. On the first 3 days, male mice were injected with saline and then placed into the recording chamber for 5 minutes before a female was added and the interaction recorded for 200 s to determine baseline number of vocalizations. On the fourth day, a drug was injected prior to the same recording procedure. Mice were given at least 3 days to recover before the start of a next 4-day protocol of a different treatment. For drug comparison, we tested 4 treatment conditions (psilocybin, 1 mg/kg, i.p.; 5-MeO-DMT, 20 mg/kg, i.p.; ketamine, 10 mg/kg, i.p.). For dose dependence, we tested 6 treatment conditions (psilocybin; 0, 0.13, 0.25, 0.5, 1, 2 mg/kg; i.p.). A randomized block design was employed such that each mouse would cycle through all treatment types. The .wav files were analyzed offline, with vocalizations detected and classified into subtypes using automated routines based on machine learning algorithms using the VocalMat software [42]. The subtypes were chevron, step up, up FM, step down, two steps, short, flat, reverse chevron, multiple steps, complex, and down FM (see **Figure 2**, below). Sounds that were detected and then subsequently classified as noise were excluded from the USV counts.

### Surgery

Surgery was performed following procedures in our prior study [31]. Briefly, mice were injected with carprofen (5 mg/kg, s.c.; 024751, Henry Schein Animal Health) and dexamethasone (3 mg/kg, i.m.; 002459, Henry Schein Animal Health) prior to surgery. Mice were anesthetized with isoflurane (3-4% for induction and 1-1.5% for maintenance) and fixed in a stereotaxic apparatus (David Kopf Instruments) with a water-circulating heating pad (Stryker Corp) set to 38°C under the body. Scalp hair was clipped, and the scalp cleaned with betadine and ethanol. Skin and connective tissue overlying the skull were removed using sterile instruments. A ∼3mm circular craniotomy was created above the right medial frontal cortex (center position: +1.5 mm anterior-posterior, AP; +0.4 mm medial-lateral, ML; relative to bregma) using a dental drill. Artificial cerebrospinal fluid (ACSF, containing (in mM): 135 NaCl, 5 HEPES, 5 KCl, 1.8 CaCl2, 1 MgCl2; pH 7.3) was used to irrigate the exposed dura above brain. A two-layer glass window was made from two round 3-mm-diameter, #1 thickness glass coverslip (64-0720 (CS-3R), Warner Instruments), bonded by UV-curing optical adhesive (NOA 61, Norland Products). The glass window was carefully placed over the craniotomy and adhesive (Henkel Loctite 454) was used to secure the glass window to the surrounding skull. A stainless steel headplate was affixed on the skull using C&B Metabond (Parkell) surrounding the glass window. Carprofen (5 mg/kg, s.c.) was given immediately after surgery and carprofen (5 mg/kg, s.c.) and dexamethasone (3 mg/kg, i.m.; 002459, Henry Schein Animal Health) were given on each of the following 3 days. Mice recovered for at least 10 days after the surgery prior to the start of imaging experiments.

### Two-photon imaging

The excitation was a Ti:Sapphire ultrafast femtosecond laser (Chameleon Ultra II, Coherent), with intensity controlled by a Pockels cell (350-80-LA-02, Conoptics) and a shutter (LS6ZM2, Uniblitz via Vincent Associates). The beam was directed into a two-photon microscope (Movable Objective Microscope) that included a water-immersion high-numerical aperture objective (XLUMPLFLN, 20X/0.95 NA, Olympus). Excitation wavelength was 920 nm and emission from 475 – 550 nm was collected using a GaAsP photomultiplier tube (H7422-40MOD, Hamamatsu). The laser power measured at the objective was ≤ 40 mW. The two-photon microscope was controlled by ScanImage 2019 software (MBF Bioscience). During imaging sessions, mice were head-fixed and anesthetized with 1-1.5% isofluorane and body temperature was controlled using a heating pad and DC Temperature Controller (40-90-8D, FHC) with rectal thermistor probe feedback. Imaging sessions did not exceed 2 hours. Apical tuft dendrites were imaged at 0 – 400 μm below the dura. For Cg1/M2, we imaged within 0 – 400 μm of the midline as demarcated by the sagittal sinus. 4 – 9 fields of view were collected from each mouse, taking 10 – 40-μm-thick image stacks at 1 μm steps at 1024 × 1024 pixels with 0.11 μm per pixel resolution. The same imaging parameters were used for every imaging session. To image the same fields of view across multiple days, we would identify and return to a landmark on the left edge of the glass window. Each mouse was imaged on days -3, -1, 1, 3, 5, 7 and 34 relative to the day of treatment. On the day of treatment (day 0) mice were injected with 5-MeO-DMT (20 mg/kg, i.p.) or vehicle. After injection, each mouse was returned to its home cage and visually observed for head twitches for 10 min.

### Analysis of the imaging data

Two-photon imaging data was analyzed using ImageJ [43] with the StackReg plug-in [44] for motion correction. If a protrusion extended for > 0.4 μm from the dendritic shaft, it was counted as a dendritic spine. The head width was measured at the widest part of the spine head using the line segment tool. Change in spine density and spine head width across imaging sessions was calculated as fold-change from the value measured on the first imaging session (day -3) for that dendritic segment. Spine formation rate was calculated as the number of new dendritic spines formed between two consecutive imaging sessions divided by the total number of dendritic spines seen in the first imaging session. Spine elimination rate was calculated as the number of dendritic spines lost between two consecutive imaging sessions divided by the total number of dendritic spines seen in the first imaging session.

### Statistics

Sample sizes were estimated based on our prior study of psilocybin [31]. GraphPad Prism 9, R, and Python [45-47] were used for statistical analyses. For head twitch response, the number and duration of head twitches between groups was analyzed by one-way ANOVA. For ultrasonic vocalizations, number of vocalizations were analyzed using Wilcoxon signed rank test. The proportions of different types of USVs were compared using the chi-square test. For in vivo two-photon imaging, dendritic spine scoring was performed blind to treatment and time. Longitudinal changes in spine density, spine head width, formation and elimination rates were analyzed using a mixed effects model for repeated measures with the *lme4* package in R. This model was chosen due to fewer assumptions being made about the underlying data (e.g., balanced sampling, compound symmetry) compared with commonly used repeated measures ANOVA. Separate mixed effects models were created for each of four dependent variables: fold-change in spine density, fold-change in average spine head width, spine formation rate, and spine elimination rate. Each model was initially run with fixed effects for treatment (5-MeO-DMT vs. vehicle), sex (female vs. male), and time (Day 1, 3, 5, and 7) as factors, including all second and higher-order interactions between terms. No sex effects were observed so final statistics reported consider only treatment and time as factors. Importantly, variation within mouse and dendrite across days was accounted for by including random effects terms for dendrites nested by mice. P-values were calculated by likelihood ratio tests of the full model with the effect in question against the model without the effect in question. Post hoc t-tests were used to contrast 5-MeO-DMT and vehicle groups per day.

### Data and code availability

The data that support the findings and the code used to analyze the data in this study will be made publicly available at https://github.com/Kwan-Lab.

## RESULTS

### 5-MeO-DMT elicits brief head-twitch response in mice independent of dose

Head-twitch response is a classic assay for characterizing psychedelics in animals [48]. The behavioral measure builds on the finding that after systemic administration of a psychedelic, the mice exhibit rapid side-to-side movements of the head. The ability of a compound to elicit head twitch response in mice correlates with its hallucinogenic potency in humans [49]. Conventionally, head-twitch responses were measured by video recording using an overhead high-speed camera (e.g., [31]). Recent studies have demonstrated that a head- or ear-mounted magnet can be used, with movements detected as electrical signals generated in a wire coil [39,40]. Here, we implemented a similar setup to use a magnetic ear tag to measure head movements in C57BL/6J mice. We refined the approach through several design iterations to finalize on a single magnetic ear tag made with a 3-mm diameter neodymium magnet (**Figure 1A**). To assess performance, we compared head twitches detected using automated procedures via magnetic ear tag versus those identified manually in video recordings by two blinded observers. The validation data set included 14 mice that received intraperitoneal injection of either saline vehicle, 1 mg/kg psilocybin, 5 mg/kg 5-MeO-DMT, or 10 mg/kg 5-MeO-DMT (**Figure 1B**). In total, 244 events were detected via magnetic ear tag versus 243 events manually identified, with 234 matched events (98.7% positive predictive value). There were 3 false positives (1.27% false discovery rate) and 2 false negatives (0.85% false negative rate). Therefore, automated detection of head-twitch response via a magnetic ear tag is highly reliable. A parts list and a step-by-step guide to construct the setup is available on a public repository (see Materials and Methods).

**Figure 1:**
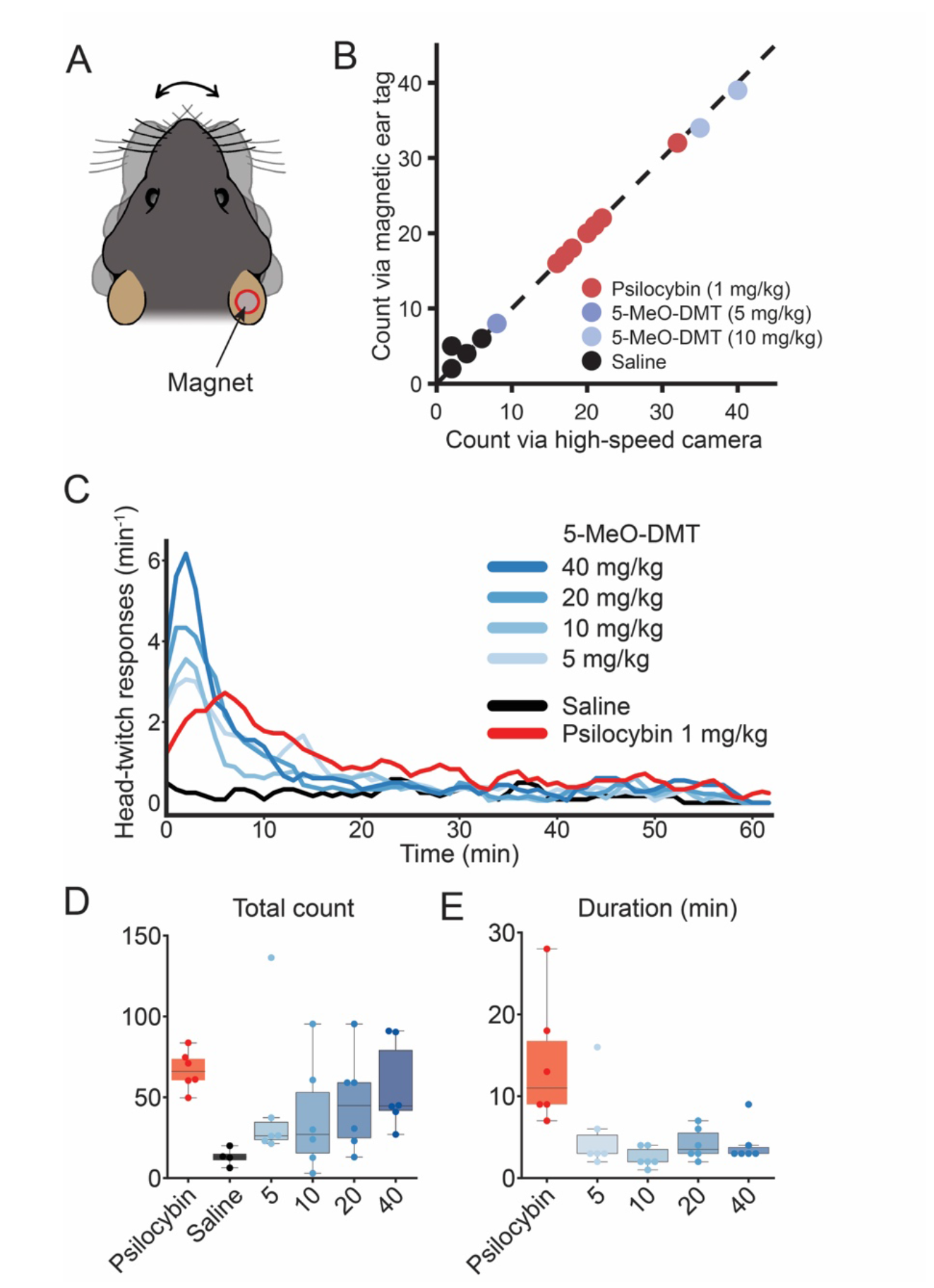
5-MeO-DMT elicits a briefer duration of head-twitch response than psilocybin. **(A)** Schematic of a C57BL/6J mouse with a magnetic ear tag. **(B)** Head-twitch response counted manually from video recordings versus automatically via the magnetic ear tag system. The Pearson correlation coefficient R^2^ = 0.996. **(C)** Head-twitch response as a function of time after systemic administration of varying doses of 5-MeO-DMT, saline, or psilocybin. **(D)** Total number of head-twitch response for 2-hr period after systemic administration. **(E)** Duration of head-twitch response, quantified by calculating the full-width half-maximum from the time course in (C). For (B), n = 14 mice. For (C - E), n = 34 mice (3 males and 3 females for each condition, except 3 males and 1 female for saline condition).

**Figure 2:**
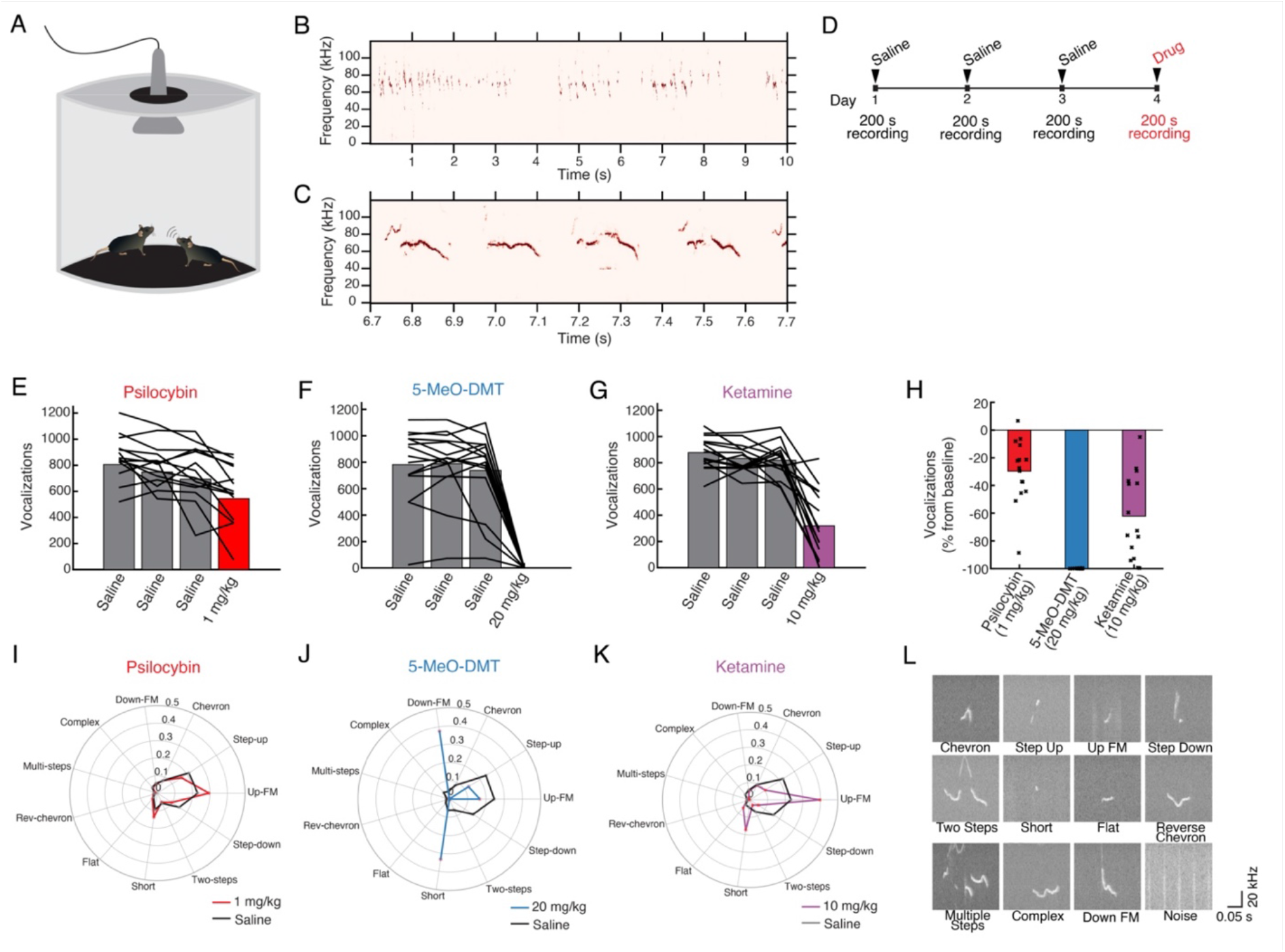
Classical psychedelics and ketamine suppress the production of social ultrasonic vocalizations. **(A)** Schematic of the setup. **(B - C)** Example recordings of USVs. **(D)** Timeline of experiment. **(E)** Number of USVs recorded on sessions before and after 1 mg/kg psilocybin administration. Bar, mean. Line, individual animals. **(F)** Similar to (E) for 20 mg/kg 5-MeO-DMT. **(G)** Similar to (E) for 10 mg/kg ketamine. **(H)** Results in (E – G) tabulated as percent changes. Bar, mean. Circle, individual animals. **(I)** Proportions of USVs classified into the 11 types, comparing sessions before and after 1 mg/kg psilocybin administration. Black, saline. Red, psilocybin. **(J)** Similar to (I) for 20 mg/kg 5-MeO-DMT. **(K)** Similar to (I) for 10 mg/kg ketamine. **(L)** Example USV recordings for each of the 11 types, as well as an example sound classified as noise. n = 15 mice.

To determine the duration of the acute action for 5-MeO-DMT, we measured head-twitch response using magnetic ear tags for 2 hr in 34 male and female C57BL/6J mice that were split into 6 groups: 5-MeO-DMT (5, 10, 20, or 40 mg/kg; i.p.), psilocybin (1 mg/kg; i.p.), and saline vehicle (equivalent volume; i.p.). As expected, psilocybin and 5-MeO-DMT elicited robust head-twitch responses (**Figure 1C**). Escalating doses of 5-MeO-DMT led to more head-twitch responses, up to the highest dose tested (**Figure 1D**). The dose-dependent increase up to 40 mg/kg is consistent with prior results that evaluated doses up to 20 mg/kg [23]. Intriguingly, animal-to-animal variability was high in the total number of head-twitch responses elicited by 5-MeO-DMT (5 mg/kg: 45 ± 18, 10 mg/kg: 38 ± 14, 20 mg/kg: 47 ± 12, 40 mg/kg: 56 ± 11, mean ± SE), relative to psilocybin (68 ± 5), echoing the varied experiences reported for 5-MeO-DMT in human studies [10,17], though some variations may have a genetic basis [50]. We quantified the duration of peak head-twitch responses by calculating full-width at half-maximum (**Figure 1E**). This analysis showed that the duration of 5-MeO-DMT’s effects was consistently brief regardless of the dose tested (5 mg/kg: 5 ± 2 min, 10 mg/kg: 3 ± 1 min, 20 mg/kg: 4 ± 1 min, 40 mg/kg: 4 ± 1 min, mean ± SE; *P* = 0.5, one-way ANOVA with 5-MeO-DMT dose as factor), relative to that of psilocybin (14 ± 3 min; any dose of 5-MeO-DMT vs. psilocybin: *P* = 0.001 treatment effect one-way ANOVA, *P* =0.002-0.03, post hoc comparisons). Previously our lab reported the behavioral and neural effects of 1 mg/kg psilocybin [31], therefore we wanted to study a dose of 5-MeO-DMT that would elicit comparable number of head-twitches. We chose to use 20 mg/kg of 5-MeO-DMT for subsequent studies. Overall, these results demonstrate the rapid duration of acute action of 5-MeO-DMT in mice.

### 5-MeO-DMT substantially reduces ultrasonic vocalization in adult mice

To further assess the acute effect of psychedelics including 5-MeO-DMT on behavior, we measured ultrasonic vocalizations (USVs) produced during mating behavior between a pair of male and female adult mice (**Figure 2A**). During such encounters, the male makes social USVs repeatedly, [41,51]; females also vocalize, but less frequently [52], so our recorded USVs likely came mostly from males (**Figure 2B**-**C**). To determine how pharmacological perturbations may impact USV production, we administered saline on 3 baseline days and then the drug of interest on the subsequent test day, recording USVs for 200 s after injection (**Figure 2D**). We compared psilocybin (1 mg/kg, i.p.), 5-MeO-DMT (20 mg/kg, i.p.), and subanesthetic ketamine (10 mg/kg, i.p.). These doses were selected because our prior studies showed that 1 mg/kg of psilocybin promotes neural plasticity [31,53], and 10 mg/kg of ketamine elevates postsynaptic calcium signaling and promotes neural plasticity [53-55]. The different drugs were tested in the same mice using a randomized block design with 7 days between successive drug administration (**Figure 2E–H**). We found that psilocybin reduced the number of social USVs (-30 ± 6%, mean ± SE; *P* = 2 × 10^−4^, Wilcoxon signed-rank test). The 5-MeO-DMT-treated mice had an even greater decrease in social USVs (-99.90 ± 0.03%; *P* = 6 × 10^−5^). However, these effects were not selective for classical psychedelics, as USV production also declined after the administration of subanesthetic ketamine (-62 ± 8%; *P* = 6 × 10^−5^).

Social USVs consist of different types characterized by distinct spectrotemporal features. In this study, USVs were detected and classified into 11 types using automated routines based on a machine learning approach via the VocalMat software [42]. The analysis revealed that the drugs not only broadly reduced the number of USVs, but also altered the mouse’s tendencies to produce certain types of USVs. Specifically, for psilocybin, mice favored the short USVs such as “up-FM” and “short”, at the expense of step sounds such as “step-up” and “step-down” (χ^2^ = 408.5, *P* < 10^−5^, chi-square test; **Figure 2I**). For 5-MeO-DMT, the near-complete suppression of social USVs precluded a reliable determination of the distribution (χ^2^ = 55.3, *P* < 10^−5^; **Figure 2J**). Ketamine produced similar effects on the proportions of USV types to psilocybin, with increased short USV types (χ^2^ = 1543.4, *P* < 10^−5^; **Figure 2K**). Prototypical examples of USVs classified as each type is shown in **Figure 2L**. Collectively, these results demonstrate that psychoactive drugs including 5-MeO-DMT can modify innate behaviors and that 5-MeO-DMT produces a profound reduction in social USV compared with more modest effects of psilocybin and ketamine.

### 5-MeO-DMT induces long-lasting increases in dendritic spine density

To measure the effects of 5-MeO-DMT on structural neural plasticity, we used longitudinal two-photon microscopy to track apical dendritic spines in the cingulate/premotor (Cg1/M2) region of the medial frontal cortex of *Thy1*^*GFP*^ mice (line M), in which a sparse subset of layer 5 and 6 pyramidal neurons express GFP [56] (**Figure 3A**). Imaging was performed for two sessions prior to injection of 20 mg/kg 5-MeO-DMT or vehicle and at two-day intervals for one week following injection, followed by one session ∼1 month later for a total of 7 imaging sessions (**Figure 3B**). We visualized the same dendrites including dendritic spines across sessions to track the number of stable, new, and eliminated spines (**Figure 3C**). A total of 870 spines were tracked across 86 dendritic segments from 10 animals (4 females, 6 males). An experimenter blind to treatment condition analyzed spine morphology according to standardized procedures [57]. To determine statistical significance of the results, we used a mixed-effects model including treatment, sex, and time as factors as well as all interaction terms. Variation within mouse and dendrite across days was accounted by including random effects terms for dendrites nested within mice. Initial analysis with the full model did not detect any sex differences, therefore the final statistical model included only treatment and time as factors. Spine density was expressed as fold-change from the first imaging session.

**Figure 3:**
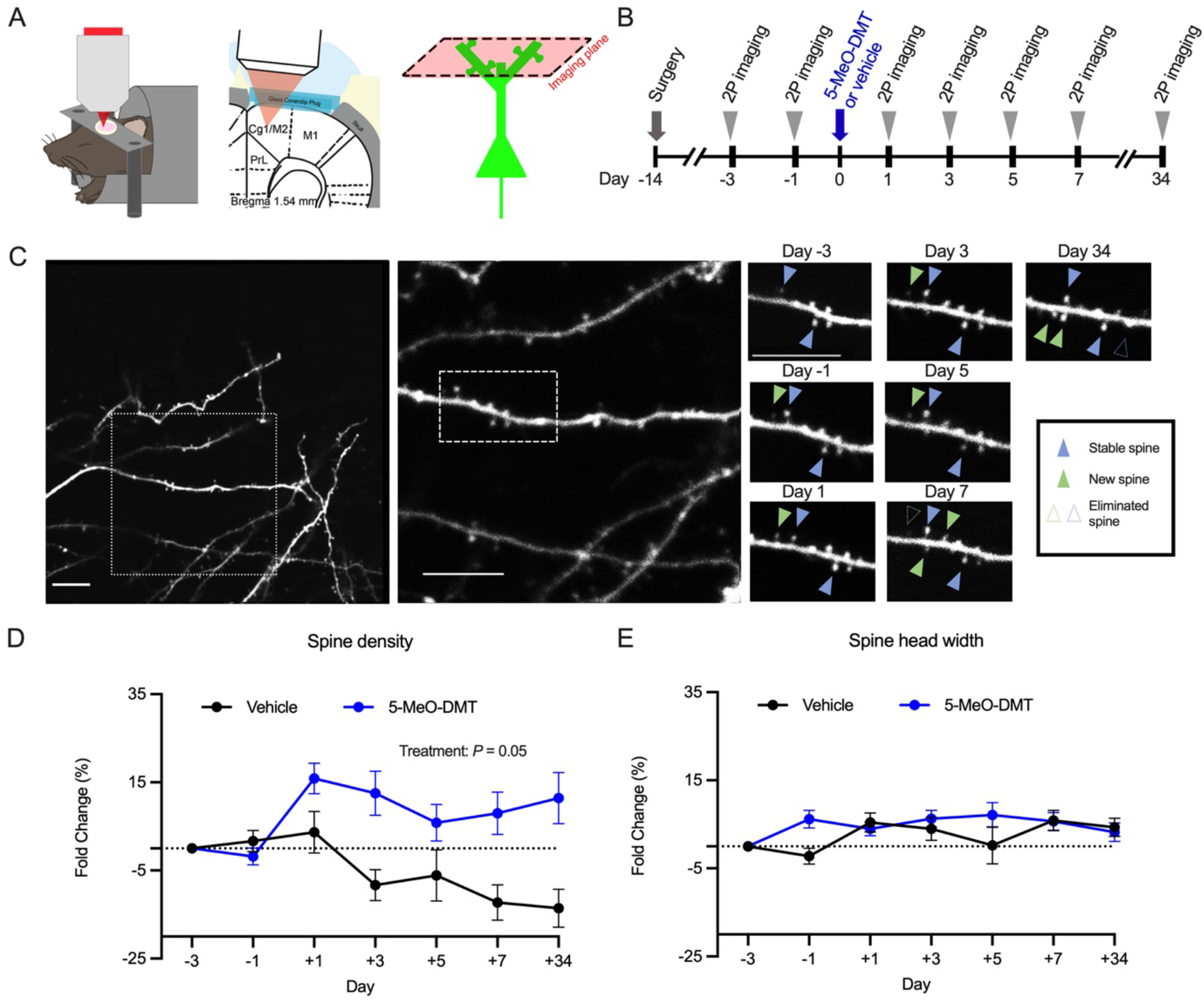
5-MeO-DMT increases the density of dendritic spines, but does not affect spine size, in the mouse medial frontal cortex. **(A)** Imaging setup. **(B)** Timeline for longitudinal imaging. **(C)** Example field of view in a mouse treated with 5-MeO-DMT. Scale bar, 5 μm. **(D)** Effects of vehicle or 5-MeO-DMT on spine density, plotted as fold-change relative to baseline Day - 3. Mean ± SEM. **(E)** Effects of vehicle or 5-MeO-DMT on spine head width, plotted as fold-change relative to baseline Day -3. Mean ± SEM. n = 6 for 5-MeO-DMT and 4 for saline

We found that a single dose of 5-MeO-DMT leads to rapid increase in spine density (5-MeO-DMT: +16 ± 3% versus vehicle: 3.6 ± 5% on Day 1, mean ± SEM), which was persistent and could be observed for at least 1 month following injection (5-MeO-DMT: +11 ± 6% versus saline: -14 ± 4% on Day 34; main treatment effect, *P* = 0.05, mixed-effects model; **Figure 3D**). We note that there was a decline in spine density observed in the vehicle group, which may due to several factors. One explanation is that a subset of the mice was treated with freebase 5-MeO-DMT (see Materials and Methods), for which the vehicle included ethanol that could have adverse effects on spine density [58]. A second reason is that the use of high laser power can damage dendritic segments, leading to declines in control groups that have been also observed in other two-photon spine imaging studies [55,59]. The ethanol and imaging conditions were matched across the two groups and therefore any extraneous effects should not affect the difference between the groups. We also measured the width of the dendritic spines, because the spine size is an indicator of the strength of the synaptic connection [60,61]. We did not detect any effect of 5-MeO-DMT on spine head width over time (main treatment effect, *P* =0.2, mixed-effects model; **Figure 3E**). Overall, these results demonstrate that 5-MeO-DMT leads to enduring increase in dendritic spine density, but did not influence the spine size in the mouse medial frontal cortex.

### 5-MeO-DMT elevates the formation rate of dendritic spines

Since we were tracking the same dendritic spines across days, we could determine the turnover rates of dendritic spines. To determine formation and elimination rates, we analyzed the same dendritic segments from consecutive imaging sessions and quantified the number of new spines formed or old spines eliminated relative to the total number of spines in the first imaging session (**Figure 4A**). We found that the formation rate increased following 5-MeO-DMT administration (5-MeO-DMT: 12 ± 1% on Day -1 and 24 ± 3% on Day 1; Saline: 7 ± 2% on Day -1 and 11 ± 3% on Day 1, mean ± SE; main treatment effect: *P* = 0.04, mixed-effects model; **Figure 4B**). This elevation in spine formation rate was transient, with post-hoc testing showing significant elevations specific to for the first and third days after 5-MeO-DMT administration (Day 1: *P* = 0.007, Day 3: *P* = 0.006). Elimination rate was not significantly affected by the 5-MeO-DMT treatment (main treatment effect: *P* = 0.96, mixed-effects model; **Figure 4C**). Altogether, these data suggest that the prolonged increase in spine density is driven by a transient increase in spine formation rate in the few days following 5-MeO-DMT administration.

**Figure 4:**
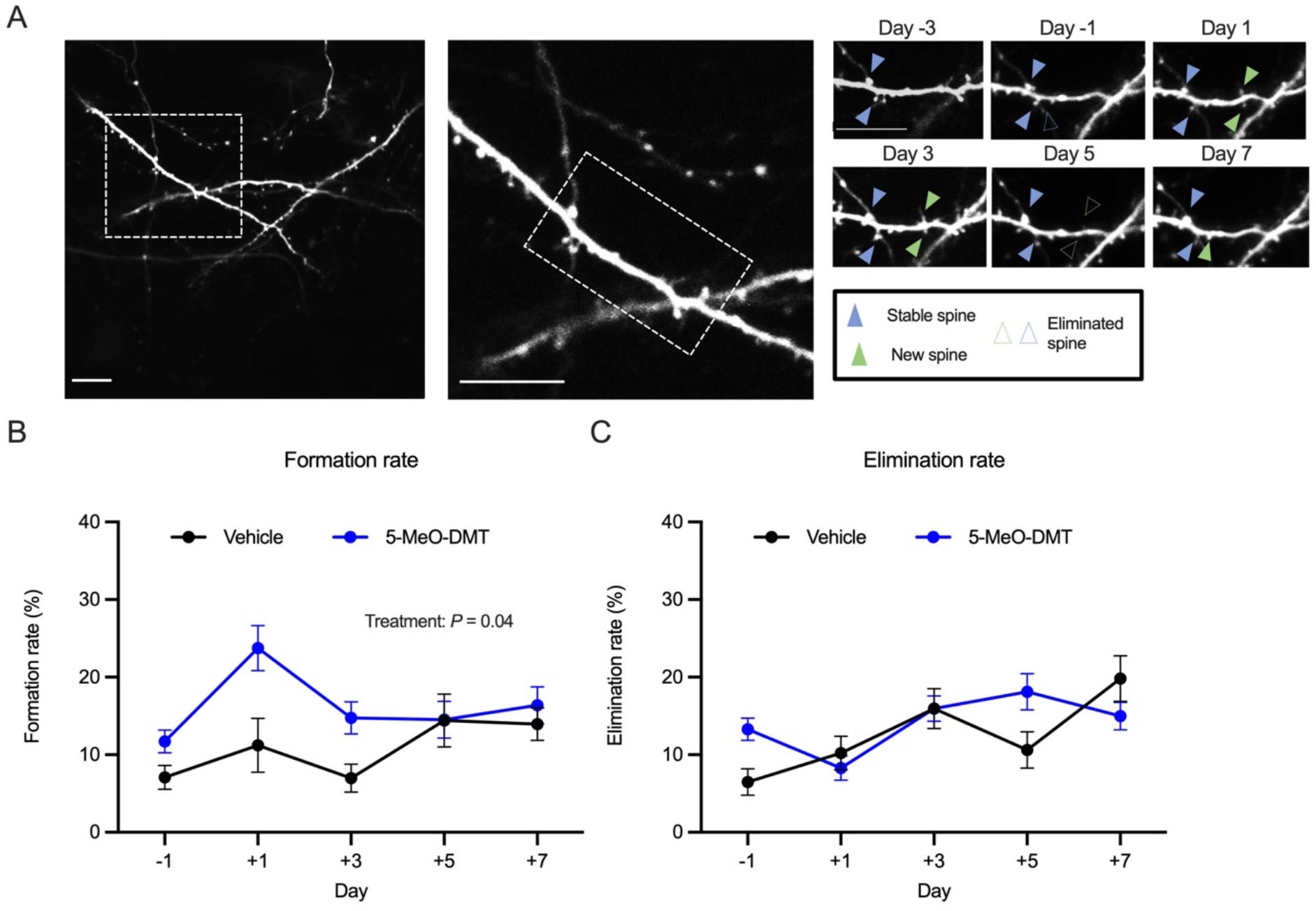
5-MeO-DMT increases the formation rate of dendritic spines. **(A)** Example field of view in a mouse treated with 5-MeO-DMT. Scale bar, 5 μm. **(B)** Effects of vehicle or 5-MeO-DMT on spine formation rate. Mean ± SEM. **(C)** Effects of vehicle or 5-MeO-DMT on spine elimination rate. Mean ± SEM. n = 6 for 5-MeO-DMT and 4 for saline

## DISCUSSION

This study yielded two main conclusions. First, 5-MeO-DMT modifies innate behaviors by increasing head-twitch response and suppressing social USVs. Notably, for head-twitch response, the effect of 5-MeO-DMT is significantly briefer than that of psilocybin at all doses tested. Second, 5-MeO-DMT induces structural plasticity in the medial frontal cortex by evoking a long-lasting increase in dendritic spine density, although intriguingly there is no effect on spine size.

5-MeO-DMT induces an extremely swift and brief subjective experience in humans due to its rapid metabolism [10,11], although the time of onset and duration can differ depending on the route of administration [12,17]. Recreational use and clinical tests typically involve smoking or inhalation of vaporized freebase, which produces a quicker onset compared to other parenteral routes such as intramuscular injection and intranasal administration [17], though all provide a more rapid onset compared to oral administration of psilocybin. For mice, vapor-based administration methods are less practical, therefore we used intraperitoneal injections. The resulting pharmacokinetics for 5-MeO-DMT in rodents, particularly in terms of the temporal profile of its entry and removal from the central nervous system, remains to be clarified. Nevertheless, behaviorally, this study confirms that when both 5-MeO-DMT and psilocybin are administered intraperitoneally, the duration of acute action for 5-MeO-DMT is considerably shorter than psilocybin at all tested doses of 5-MeO-DMT. A previous study has shown that the head-twitch response elicited by the administration of 5-MeO-DMT is dependent on 5-HT_2A_ receptor agonism [23], like psilocybin.

Given the paucity of information on how psychedelics impact innate behavior beyond head-twitch response, we decided to also examine the impact of various compounds on social USV. In the presence of females, male mice spontaneously produce USVs [41,51], which are frequent and have quantifiable spectrotemporal features, making this an attractive behavioral paradigm. Moreover, studies have delineated the neural circuit involved in vocal control, which include premotor neurons in brainstem and periaqueductal grey that are controlled by cortical regions including motor cortex and medial frontal cortex [62]. The behavioral assay is relevant because the medial frontal cortex is the location where current and past studies have shown psychedelic-evoked plasticity [31,36,37]. The periaqueductal grey, which has been shown to be essential for social USVs in mice [63], is impacted by serotonergic neuromodulation [64], and by psychedelics [53]. Here we demonstrate that psilocybin and 5-MeO-DMT robustly suppress the production of social USVs and alter the pattern of USVs produced. Ketamine has a similar effect. The results could be due to effects of psychedelics on motor control of vocal production, given that 5-HT_1A_ agonists and NMDAR antagonists have been reported to impair distress vocalizations in maternally separated mouse pups [65,66]. Alternatively, the results could be due to effects of psychedelics on social behavior. Interestingly, the most profound effects on social USV were seen with 5-MeO-DMT, suggesting that suppression of USV may correlate with intensity of psychedelic effects. Social USV may be a useful assay for assessing the acute effect of psychedelics in animal models.

It is worthwhile to compare the effects of 5-MeO-DMT and psilocybin on structural plasticity. We intentionally designed the current experiment to match the timeline used in our prior study [31], in order to facilitate a direct comparison. Relative to psilocybin, 5-MeO-DMT evokes a similar ∼10-15% increase in spine density that rises within 1 day after administration and endures for at least 1 month. Moreover, just like psilocybin, the increase in spine density is driven by a transient increase in spine formation rate within the initial 1 – 3 days after injection. These results suggest that the effects of a single dose of 5-MeO-DMT on spine density in medial frontal cortex is indistinguishable from those reported previously for psilocybin. The notable difference is in the effects of the compounds on spine head width. Whereas psilocybin causes a ∼10% increase in spine size that sustains for days before returning to baseline within a month, 5-MeO-DMT has no detectable impact on spine head width. Spine size has been shown to have a linear relationship with synapse strength [61], so one may interpret that the average strength of synapses in the medial frontal cortex is not affected by 5-MeO-DMT on the timescale measured despite the overall number of dendritic spines being increased. It remains to be seen how these different measures of plasticity track with the therapeutic effects of these drugs.

The results have implications for drug discovery. There is a concerted push in the field to develop non-hallucinogenic psychedelic analogs [37,67,68]. The potential therapeutic effects of these new chemical entities have so far been demonstrated using behavioral assays such as forced swim, tail suspension, and sucrose preference tests. The current study provides insight into the question of whether duration of psychedelic effects correlates with duration of therapeutic effects. By showing that the short-acting 5-MeO-DMT can produce long-lasting effects on neural structural plasticity *in vivo*, the results suggest that the durations of acute and long-term effects may not be tightly linked. To conclude, 5-MeO-DMT is a compound with increasing clinical relevance. Further characterizations of its effects, particularly in contrast to psilocybin and other psychedelic analogs, will shed light onto the mechanisms of actions that support its therapeutic potential for treating mental illnesses.

## ACKNOWLEDGMENTS

We thank Ling-Xiao Shao for advice on imaging, Neil Savalia for advice on statistical analyses, Gregg Castellucci and Katherine Tschida for advice on measuring ultrasonic vocalizations, Antonio Fonseca for advice on the VocalMat software, and Janghoo Lim for loaning equipment for recording ultrasonic vocalizations. 5-MeO-DMT-succinate was generously provided by Usona Institute’s Investigational Drug & Material Supply Program; the Usona Institute IDMSP is supported by Alexander Sherwood, Robert Kargbo, and Kristi Kaylo in Madison, WI. The behavioral experiments were supported by the Yale Program in Psychedelic Science, NIH/NIMH grant R01MH121848 (A.C.K.), NIH/NIMH grant R01MH128217 (A.C.K.), One Mind – COMPASS Rising Star Award (A.C.K.), and NIH/NIGMS medical scientist training grant T32GM007205 (P.A.D.). The imaging experiments were supported by a sponsored research project from Freedom Biosciences (C.P., S.J.J.), NIH/NINDS training grant T32NS041228 (C.L.), China Scholarship Council-Yale World Scholars Fellowship (H.W.), and NIH/NIMH training grant T32MH019961 (S.J.J.).

## AUTHOR CONTRIBUTIONS

S.J.S., C.P., and A.C.K. designed the research. S.J.S. performed the two-photon imaging experiments and analyzed the data. I.G. performed the ultrasonic vocalization experiments and analyzed the data. M.D. performed the head-twitch response experiments and analyzed the data. C.L. and H.W. assisted in the two-photon imaging experiments. P.A.D. worked with M.D. to make the equipment for using magnetic ear tags to measure head-twitch response and analyzed the data. J.S.S. contributed to the experimental design. A.M.S. synthesized and provided the 5-MeO-DMT succinate. C.P. and A.P.K. contributed to data interpretation. S.J.S. and A.C.K. wrote the paper with input from other authors.

## DISCLOSURES

A.C.K. is a member of the Scientific Advisory Board of Empyrean Neuroscience and Freedom Biosciences. A.C.K. has consulted for Biohaven Pharmaceuticals. No-cost compounds were provided to A.C.K. for research by Usona Institute. C.P. has served as a consultant in the past year for Biohaven Pharmaceuticals, Teva Pharmaceuticals, and Brainsway Therapeutics and has performed research under contract with Biohaven and with Blackthorn Therapeutics, Ltd., on unrelated projects. C.P. and A.P.K have performed research under contract with Transcend Therapeutics on unrelated projects. C.P. and S.J.S. have a sponsored research agreement with Freedom Biosciences. The sponsor of this research was not involved in the analyses or writing of the manuscript. The duties had no influence on the content of this article.

